# Pan-genome analysis of *Mycobacterium africanum*: insights to dynamics and evolution

**DOI:** 10.1101/2020.03.17.995167

**Authors:** Idowu B. Olawoye, Simon D.W. Frost, Christian T. Happi

## Abstract

*Mycobacterium tuberculosis* complex (MTBC) consists of seven major lineages with three of them reported to circulate within West Africa: lineage 5 (West African 1) and lineage 6 (West African 2) which are geographically restricted to West Africa and lineage 4 (Euro-American lineage) which is found globally. It is unclear why the West African lineages are not found elsewhere; some hypotheses suggest that it could either be harboured by an animal reservoir which is restricted to West Africa, or strain preference for hosts of West African ethnicity, or inability to compete with other lineages in other locations.

We tested the hypothesis that *M. africanum* (MAF) might have emigrated out of West Africa but was outcompeted by more virulent strains of *M. tuberculosis* (MTB).

Whole genome sequences of MTB from Nigeria (n=21), China (n=21) and MAF from Mali (n=24) were retrieved, and a pan-genome analysis was performed after fully annotating these genomes. The outcome of this analysis shows that Lineages 4, 5 and 6 have relatively close pan-genomes whilst lineage 2 has an open pan-genome. We also see a correlation in numbers of some multiple copy core genes and amino acid substitution with lineage specificity that may have contributed to geographical distribution of these lineages.

The findings in this study provides a perspective to one of the hypotheses that *M. africanum* might find it difficult to compete against the more modern lineages outside West Africa hence its localization to the geographical region.

## Introduction

Tuberculosis (TB) is the world leading cause of death from infectious diseases, with the latest TB report stating over 1.1 million deaths globally in 2018, with two thirds from low-to-middle income countries such as China, Indonesia, Nigeria, the Philippines, Pakistan, South Africa and Bangladesh (1). *Mycobacterium tuberculosis* complex (MTBC), the causative organism for tuberculosis has been studied widely and is known to be a monoclonal bacteria compared to other bacterial models acquiring variation predominantly through mutations (2). MTBC is grouped in two major ancestries, the ancient lineages and the modern lineages, both of which are further grouped into lineages and sub lineages based on the geographical regions that they are found. Lineage 1 (East Africa, Philippines, in the region of the Indian Ocean), lineage 2 (East Asia), lineage 3 (East Africa, Central Asia), lineage 4 (Europe, America and Africa), lineage 5 (West Africa 1), lineage 6 (West Africa 2) and lineage 7 in Ethiopia (3). The ancestry of these lineages was established by the study of 20 variable genomic regions that are caused by insertion or deletion events (4). For example, the absence or presence of an MTB specific deletion known as *TbD1* and regions of difference (RD) helps to classify MTBC into modern (lineages 2,3 and 4) or ancient (lineages 1, 5. 6 and 7) strains.

Different lineages of MTBC have shown to present different symptoms and immunological responses as seen in several studies. Although there is limited knowledge of TB virulence, pathogenicity does not associate with classical virulence factors like toxins (5), but rather with other complex factors, such as other bacterial infections being harboured by the patient that can interact with the tubercle bacilli, the host immune system, or even environmental factors resulting in a complex cascade of events (6). Furthermore, reports have linked TB infectivity to human genetic susceptibility, with certain polymorphisms in the human genome relating to certain MTBC lineages in their respective geographic distribution (7–9). Interestingly, a study found that a particular variant in a gene responsible for autophagy in humans, IRGM-261T, influences the susceptibility of TB caused by MAF but not MTB (10). Additionally, findings from research on Ghanian populations showed that variants associated with 5-lipoxygenase (ALOX5) waere associated with an increased TB risk (11). This suggests that MTBC might be adapted to certain populations just as they are geographically distributed (12).

In addition to host-pathogen compatibility of TB, environmental factors, lifestyle, living conditions and HIV co-infection play important roles in the outcome of the infection (13).

In Nigeria and West Africa at large, only three major lineages have been reported to cause tuberculosis: lineage 4, also known as the Euro-American lineage and lineages 5 and 6 also known as West Africa lineages 1 and 2. Lineage 4 comprises of 10 sub lineages: L4.1.1 (X); L4.1.2 (Haarlem); L4.1.3 (Ghana); L4.2; L4.3 (LAM); L4.4; L4.5; L4.6.1 (Uganda); L4.6.2 (Cameroon); and L4.10 (PGG3) (14). Lineage 4 is globally distributed, with the majority of TB infections in Nigeria bing caused by the Cameroon sub lineage of lineage 4, followed by *M. africanum* (Lineages 5 and 6) (15).

With the advent of next generation sequencing and the continuous reduction in the cost to sequence the entire genome of an organism, studies have moved from the analysis of a single or few genomes to multiple or a collection of genomes. Pan-genome analysis is a product of the breakthrough of multi-genome study in molecular biology (16).

The pan-genome can be described as the collection of entire genes in a particular species. This comprises the core genes shared by all strains, dispensable genes shared by two or more strains, and unique genes also known as singletons that are peculiar to specific strains. This can be used in describing bacterial species as many species differ by their gene content to a large extent (17). The core genes are responsible for the major phenotype and basic biological processes of the bacteria, whilst the accessory and unique genes may be involved in other metabolic pathways such as adaptation to a particular host, virulence, antibiotic resistance and other functions that confer selective benefits over other species (16). Pan-genome analysis has been performed on more than 50 bacteria species in the past decade and this has revealed interesting information relating to pathogenesis, bacteria evolution, drug resistance, host specialization, horizontal gene transfer (HGT) (18,19). Pan-genome analysis has also been adopted for viruses, fungi and plants (20,21). Studies have also used pan-genome analysis for identifying potential vaccine candidates against bacterial infections (18)

The purpose of this study was to compare the entire gene set of *M. africanum* and the most commonly found TB sub lineage in Nigeria L4.6.2 (also known as the Cameroon sub lineage) and L2.2.1 (also known as the Beijing lineage) as an out-group to understand the evolution, genome dynamics, metabolic pathways and also genes involved in biological processes.

## Methodology

### Sample collection and filtering

We first retrieved fully assembled genomes of *M. africanum* (n=24) out of 30 available genomes on the NCBI Reference Sequence Database (RefSeq), selected sequences were those that were not derived from surveillance projects or contain anomalies. We retrieved an additional 30 raw sequence datasets from the Senghore *et al.* study (15), and selected 21 genomes after filtering and profiling them according to respective lineages using TB-Profiler version 2.3 (22). Thus, we selected only the Cameroon sub lineage of the Euro-American lineage. An out-group set of lineages was used for comparison by obtaining 43 genome assemblies from NCBI RefSeq that were sequenced from China. These selected genomes were classified into subtype lineages using Biohansel (23), and only genomes that belongs to the Beijing sub lineage (n=21) were selected.

### Bioinformatics analysis

The genomes of the Cameroon sub lineages were assembled using a de novo approach with SPAdes version 3.11.1 (24). The scaffolds of the MAF genomes from Mali, Cameroon sub lineage and the Beijing lineage were annotated using Prokka version 1.12 (25). The annotated genomes were analysed with the Bacterial Pan Genome Analysis Tool (BPGA) version 1.3 (16), using 95% similarity for orthologous clustering and both pan- and core genomes were calculated over 20 permutations to prevent bias. Additionally, the pan-genome functional analysis was carried out utilizing KEGG, COG metabolic and functional pathways, which were all visualised with LibreOffice Calc plot functions. Panaroo (26) and PopPUNK (27) were also employed for pan-genome investigation of gene profiles and clustering core and accessory genomes.

## Results

### Mycobacterium africanum

The identification of core genes (genes shared between all strains), accessory genes (genes shared by two or more strains but not all) and unique genes (genes peculiar to individual strains) were clustered respectively. The pan-genome analysis across 24 genomes of *M. africanum* showed that they have 3974 ± 20 genes (mean ± standard deviation). The number of accessory genes ranged from 194 to 222 genes and the unique set of genes varied from 1 to 38 genes (Supplementary table 1A). Empirical power law equations and exponential equations were used to generate pan- and core genome curves, which showed that the pan-genome curve has almost reached a plateau as the exponent parameter calculated is 0.03 (Fig 1). This argues that the 24 genomes analysed are sufficient to obtain an accurate estimate of pan- and core genome size, with additional samples yielding diminishing returns.

**Fig 1:**
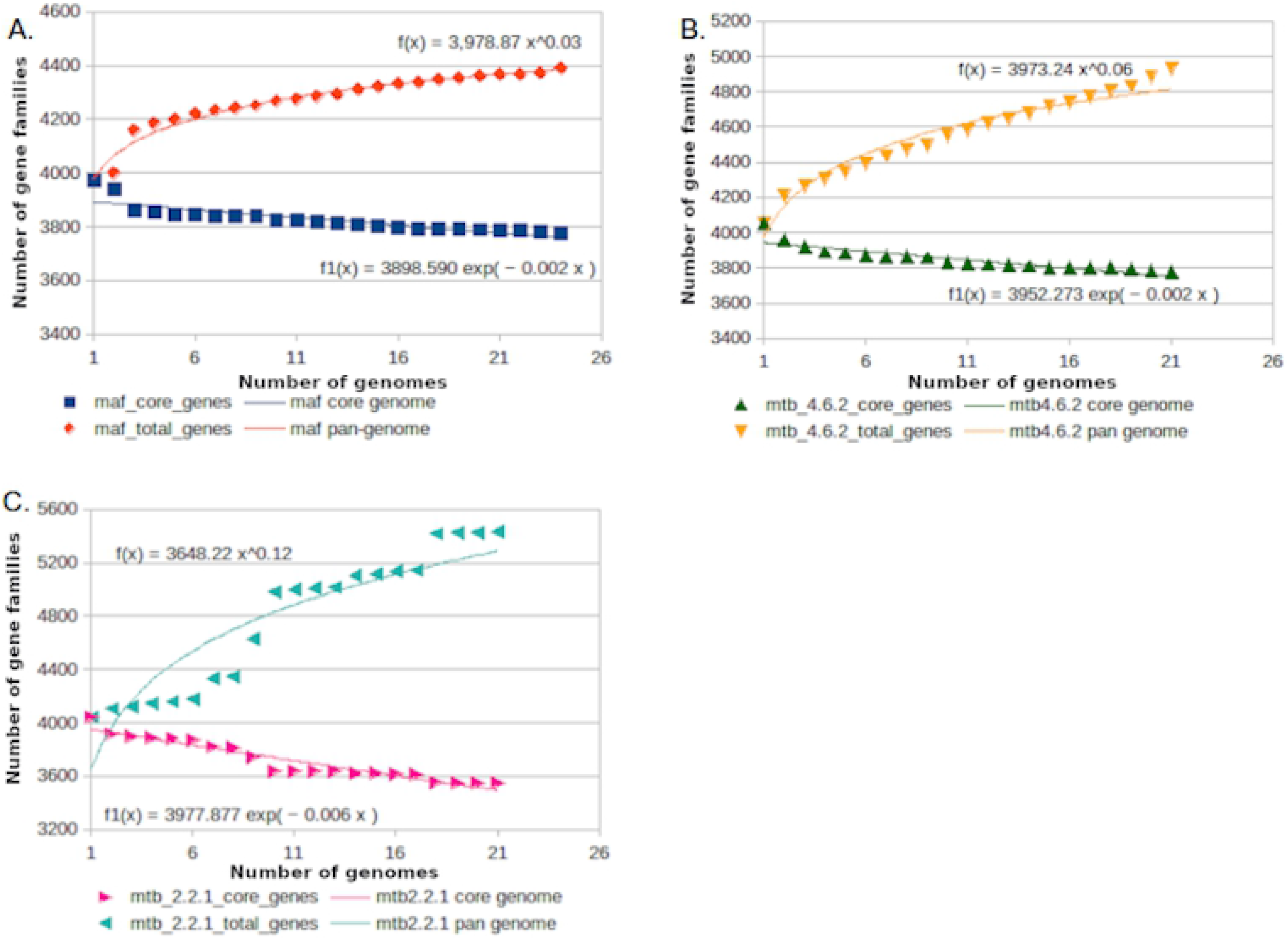
The pan and core genome of the MTBC genomes. (A) MAF: *M. africanum*, (B) MTB-4.6.2: Cameroon sub lineage and (C) MTB-2.2.1: Beijing lineage profile plot with power-fit curve equation shown as f and exponential curve equation as f1. Pan-genome functional analysis was done using KEGG pathway database and KEGG IDs assigned to orthologous protein clusters in the core, accessory and singleton genes and matched against the database, as shown in Fig 2. The highest gene contents in the pan-genome of *M. africanum* were responsible for biological processes such as metabolism, a remarkable amount of unique genes account for environmental information processing and organismal systems. A more detailed KEGG classification showed that majority of unique gene sets were responsible for signalling molecules and interaction, signal transduction, infectious diseases, digestive system and cellular community (Fig 3).

### *Mycobacterium tuberculosis* sub lineage 4.6.2 (Cameroon sub lineage)

The classification of core, accessory and unique genes in numbers after clustering into orthologous groups was performed. The 21 genomes from the Cameroon sub lineage had 4067 ± 18 genes (mean ± standard deviation). Accessory gene numbers ranged from 242 to 289 and singleton gene numbers were from 19 to 55 genes (Supplementary table 1B). Power law equations and exponential equations were used to generate pan and core genome curves which reflects the curve has almost reached a plateau as the exponent parameter calculated is 0.06 (Fig 1).

A high percentage of genes in the core genome were linked to metabolic processes after functional classification of the pan-genome (Fig 2). A more detailed KEGG classification of the pan genome showed that majority of the accessory genes are related to infectious diseases and cellular community (Fig 3).

**Fig 2:**
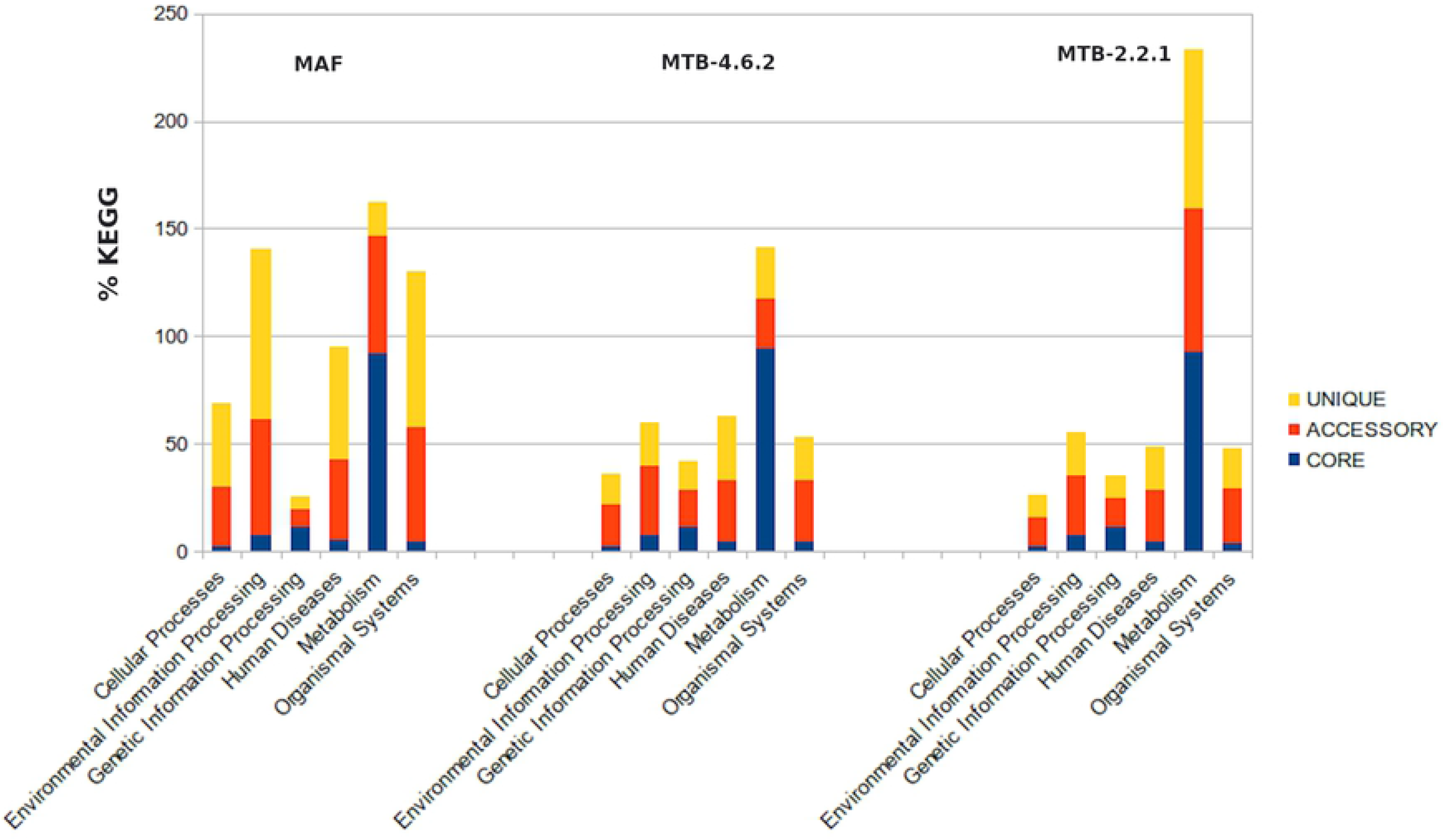
KEGG distribution of core, accessory and unique genes for the MTBC genomes. MAF: *M. africanum*, MTB-4.6.2: Cameroon sub lineage and MTB-2.2.1: Beijing lineage

**Fig 3:**
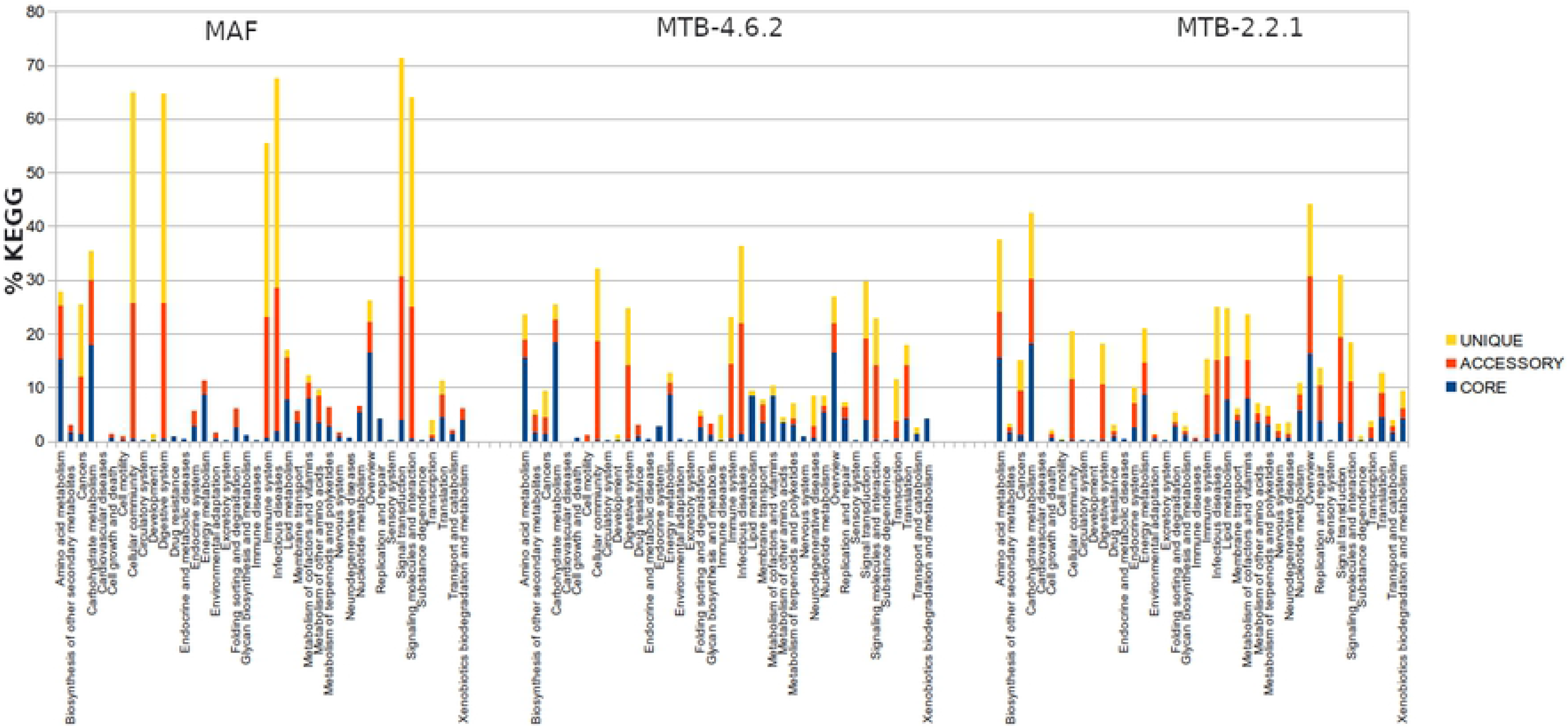
Detailed KEGG classification of core, accessory and unique genes for the MTBC genomes. MAF: *M. africanum*, MTB-4.6.2: Cameroon sub lineage and MTB-2.2.1: Beijing lineage

### *Mycobacterium tuberculosis* sub lineage 2.2.1 (Beijing sub lineage)

Orthologous clustering of the pan-genome showed that the analysis of 20 genomes have 4023 ± 83 genes (mean ± standard deviation). The number of accessory genes ranged from 359 to 435 genes, with 3546 genes shared between all strains (core genes) and unique gene sets varied from 0 to 338 genes per strain (Supplementary table 1C). Exponential regression was used to generate the number of core genes whilst power law regression was used to extrapolate the pan-genome curve which reflects an open genome with an exponent parameter 0.12 (Fig 1).

KEGG classification of core, accessory and unique genes into functional roles by clustering genes into orthologous groups argued that the majority of core and accessory genes are responsible for metabolic functions (Fig 2) and a detailed classification of functional genes showed that a relative quantity of the core genes are associated with carbohydrate metabolism, whilst the accessory genes are linked with signal transduction and infectious diseases (Fig 3).

### Pan-genome comparative analysis

Following the clustering of respective TB datasets into core, accessory and unique genes, the core genes of MAF were almost equal to the number of core genes in the Cameroon sub lineage (MTB-4.6.2) but had fewer accessory genes than the Cameroon sub lineages. However, the Beijing sub lineages (MTB-2.2.1) had fewer core genes than MAF and MTB-4.6.2, but more accessory genes than both (Supplementary table 1). The red, orange and cyan lines in Figure 1 (a, b and c) represent the power-fit curve derived from the equation (*f(x)=a.x^b*), where the exponent *b* > 0 implying that the genome is open, however the parameter *b* values are 0.03, 0.06 and 0.12 (MAF, MTB-4.6.1 and MTB-2.2.1 respectively) meaning the pan genome is almost closed and addition of new genomes may not lead to the discovery of novel functions (28).

Using BPGA to cluster the core, accessory and unique genes and assigning KEGG functional pathways to them, about 90% of the core genes of the three lineages are responsible for metabolism related pathways. However, accessory gene assignment of metabolic, organismal system and environmental information related pathways in the lineages are: MAF (54%, 52% and 53%), MTB-4.6.2 (22%, 28% and 32%) and MTB-2.2.1 (66%, 25% and 27%) as shown in Figure 2.

### Evolution of *M. africanum*

Core genes are responsible for survival and majority of biological processes in the bacteria. Those that exist in copies, also known as “multi copy core genes” (MCG) were studied as these genes, especially rRNAs in bacteria, have been seen to influence adaptation to environmental pressures and the structure of microbiomes (29). We examined conserved genes of these lineages that exhibit copy number variation (CNV) and constructed a phylogeny based on their core genomes (Figure 4). Eleven (11) Proline-Glutamate (PE)/Proline-Proline-Glutamate (PPE) family proteins and four (4) *fadD* genes fall under the category of MCG. PPE family proteins and *fadD* genes were selected as they have been reported to be crucial factors for mycobacterial virulence and *in vivo* pathogenicity (30–32).

**Fig 4:**
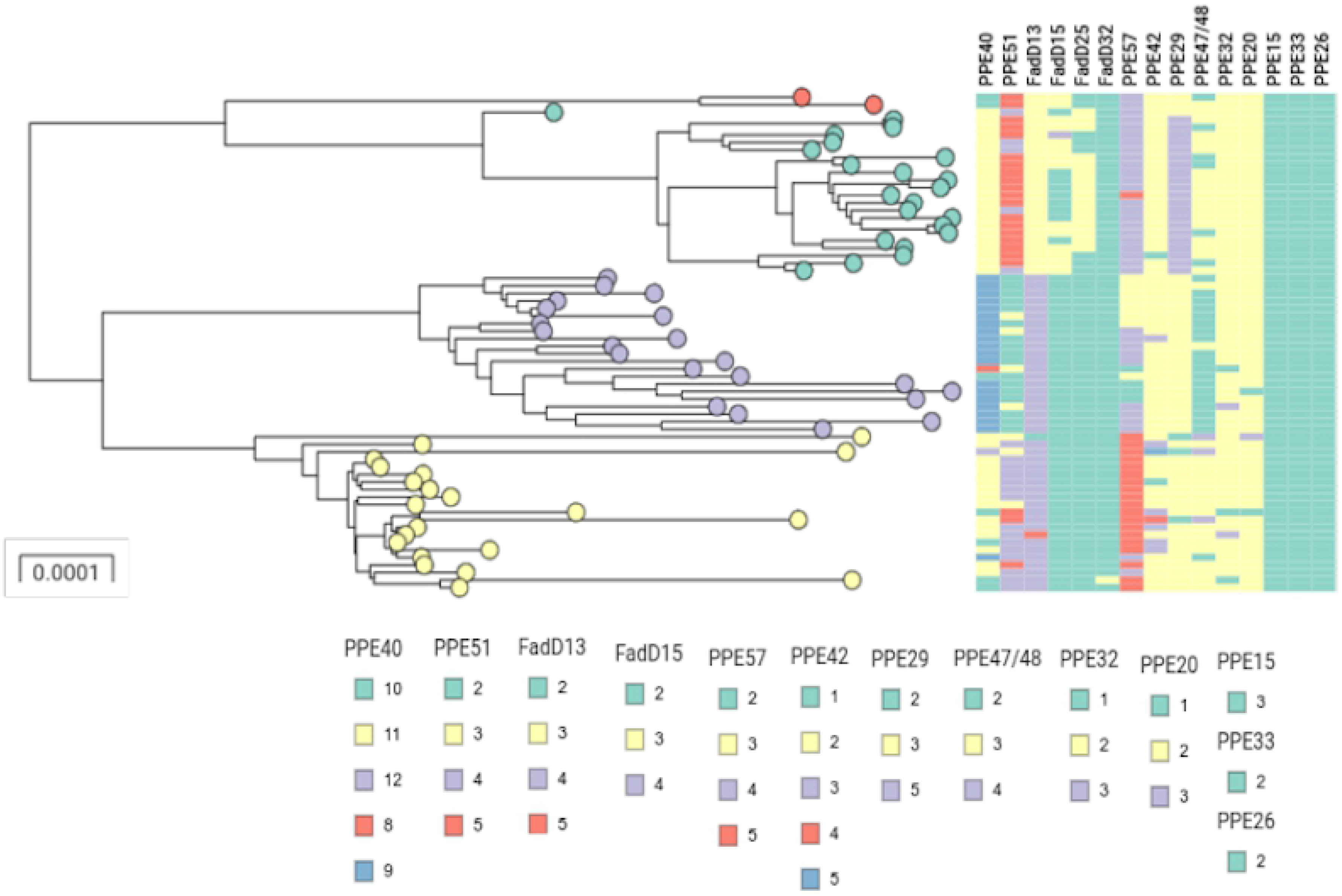
Core genome phylogeny of strains studied in this paper. Tree nodes are coloured according to clusters generated by PopPUNK; red and blue (MAF), purple (MTB-4.6.2) and yellow (MTB-2.2.1). Columns display multi-copy core PPE and *fadD* genes. Genes are coloured according to copy numbers.

PPE family proteins 15, 26 and 33 all had similar gene copy number across all strains, which shows that these proteins have remained unchanged over time, whilst family PPE proteins 20, 32 and 42 showed little evolution in the CNV and lastly, PPE proteins 40, 51 and 57 displayed lineage distinct CNV across the genomes (Fig 4).

On the other set of gene families studied, *fadD13*, *fadD15*, *fadD25* and *fadD32* genes which are responsible for fatty acid CoA ligase, all showed CNV in the TB strains during the course of evolution except the *fadD32* gene (Fig 4). Additionally, multiple sequence alignment of *fadD13* genes in MAF and MTB genomes showed substitution mutations *A8G* in *fadD13_2*, *S108F* and *A123V* in *fadD13_3* (Fig 5).

**Fig 5:**
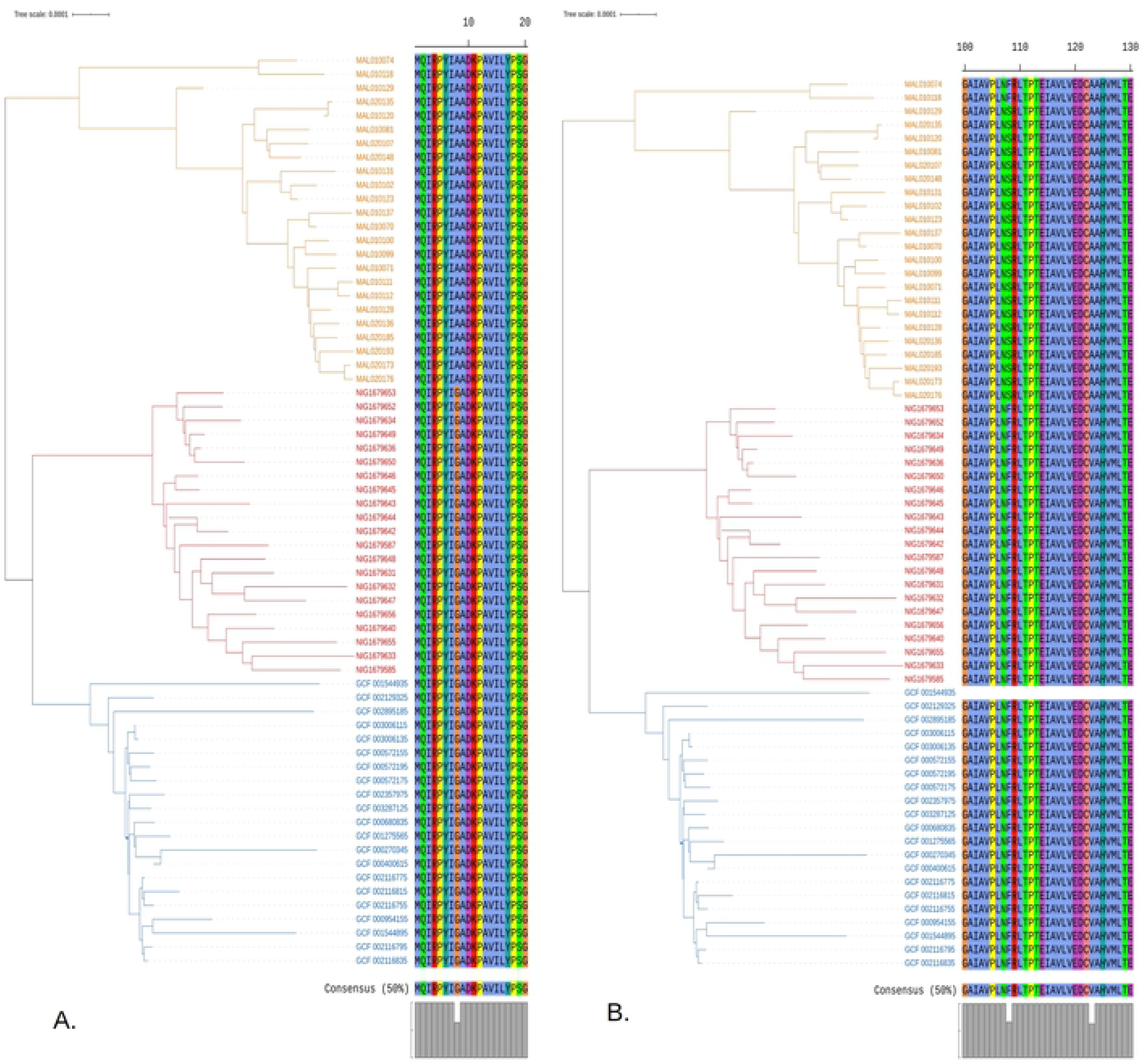
Core genome phylogeny and multiple sequence alignment of some multi copy core genes. (A) *fadD13_2* and (B) *fadD13_3* showing amino acid substitutions within lineages MAF (orange), MTB-4.6.2 (red) and MTB-2.2.1 (blue). *FadD13_1* is not included as there is no mutation in the gene. Also, *fadD13_3* is absent in sample GCF 001544935 hence the blank space.

## Discussion

Despite the clonal nature of MTBC, our findings shows that the addition of new genomes to the analysis will not lead to the discovery of new phenotypes in MAF due to its close pan-genome. In addition, we also observed copy number variations of some core genes that may be related to the geographically restricted specialists (MAF) and globally distributed generalists (MTB-4.6.2 and MTB-2.2.1).

The reduction in number of core genes in MTB-2.2.1 compared to MAF and MTB-4.6.2 can be linked to increased virulence as previous work have shown that the Beijing sub lineage is more pathogenic than ancient lineages. Genome reduction causes bacteria to have increased virulence compared to their counterparts with larger genomes (33,34). Varying numbers of accessory and unique genes which could also be responsible for drug resistance, virulence or preference to a particular host which plays a key role in lineage classification, geographical distribution and distinct lifestyles of some TB strains as reported in earlier studies (12,16). Furthermore, investigation into some MCG show copy number variation and amino acid substitution during the evolution of MTBC, this could also attribute to host preference and the geographical peculiarity of MAF.

In the detailed KEGG classification shown in Fig 3, a large number of orthologous genes belonging to cellular community, digestive system, immune system, infectious diseases, signal transduction and signalling molecule interactions pathways were grouped under unique and accessory genes in MAF, whilst a significant reduction of orthologous genes in these same pathways were seen in MTB-4.6.2 and MTB-2.2.1. We speculate that genetic loss in the dispensable genome of the modern lineages has a selective advantage for virulence and global distribution that allows the TB strain to persist in a wide range of host as reported in previous works (35,36). We also saw a reduction in copy number of some core genes in MTB-4.6.2 and MTB-2.2.1, which was higher in MAF such as PPE family protein 40, 51, *fadD13* and 15 (Fig 4). This may also be related to the geographic distribution and host specificity as housekeeping genes are responsible for survival of pathogens (37).

One of the multiple copy core genes, *fadD13*, which codes for long chain fatty acid COA ligase in MTB and maintains appropriate mycolic acid composition and permeability of the envelope on its exposure to acidic pH (38,39), was investigated for evolution in the lineages. The *fadD13* gene is encoded by operon *MymA* and is essential for virulence and *in vivo* progression of MTBC (40). Multiple alignments of gene copies of *fadD13* (Fig 5) in the TB genomes suggests that these lineage-specific mutations could have also shaped the distinctive features of the modern and ancient lineages in this study, as numerous *fadD13* protein variants have been seen to influence ATP binding (40).

## Conclusion

Pan-genome analysis of MAF in this work showed that a higher number of orthologous genes in the dispensable genome may have contributed to the restriction of global distribution and reduced virulence compared to modern lineages MTB-4.6.2 and MTB-2.2.1. Also, substitutions in some multiple copy core genes and copy number variations might have influenced evolution of MAF and its geographic distribution.

Inasmuch as it is hard to say if MAF migrated out of West Africa and got outcompeted by modern lineages, it is almost certain that it might soon be outcompeted by modern virulent strains in West Africa due to its close pan genome.

## Acknowledgement

We would like to show appreciation to our colleagues at the African Centre of Excellence for Genomics of Infectious Diseases and the facility made available at the research centre which continues to provide an enabling environment for groundbreaking research. This work is supported by grants from the National Institute of Allergy and Infectious Diseases. NIH-H3Africa (U01HG007480 and U54HG007480 to Redeemer's University [Dr. Happi]), and a grant from the World Bank grant (project ACE019 to Redeemer's University [Dr. Happi]).

## Supporting information

**Supplementary table 1: Number of core, accessory and unique genes from individual genomes** (A) MAF: *M. africanum* pan-genome, (B) MTB-4.6.2: Cameroon sub lineage pan-genome and (C) MTB-2.2.1: Beijing lineage pan-genome

